## Supplementary material for "Massively parallel identification of causal variants underlying gene expression differences in a yeast cross": All supplementary figures

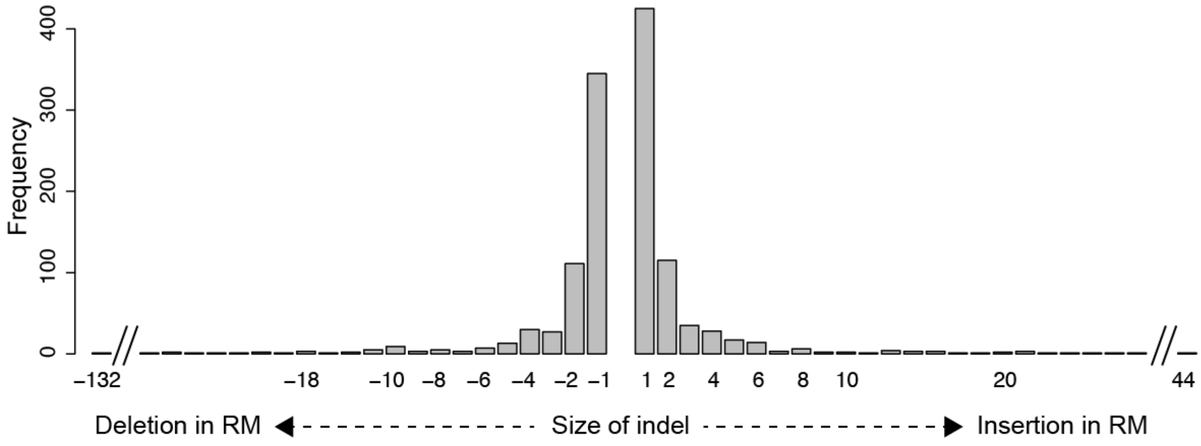


Figure S1: Size distribution of indels in the MPRA design

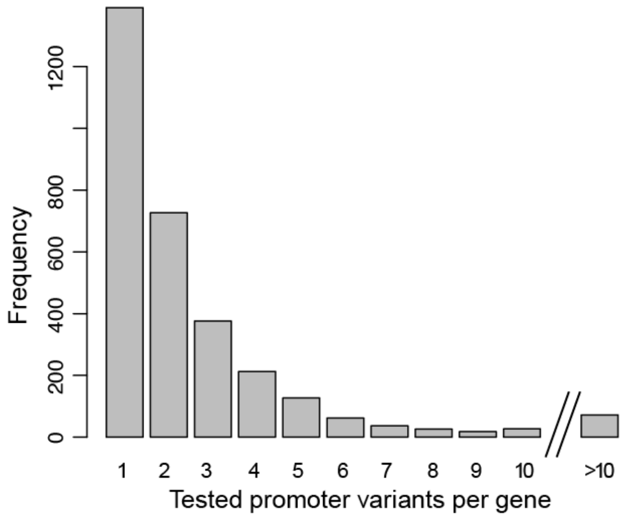


Figure S2: Distribution of the number of promoter variants per gene in the MPRA design

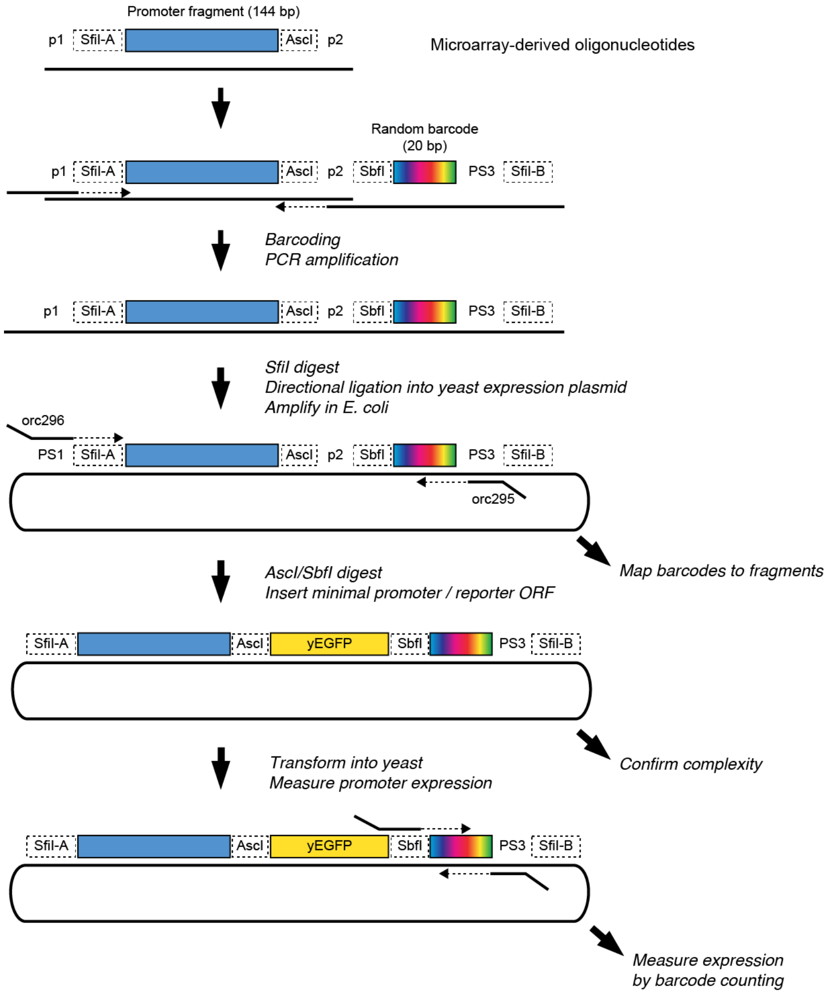


Figure S3: Schematic of the library cloning procedure.

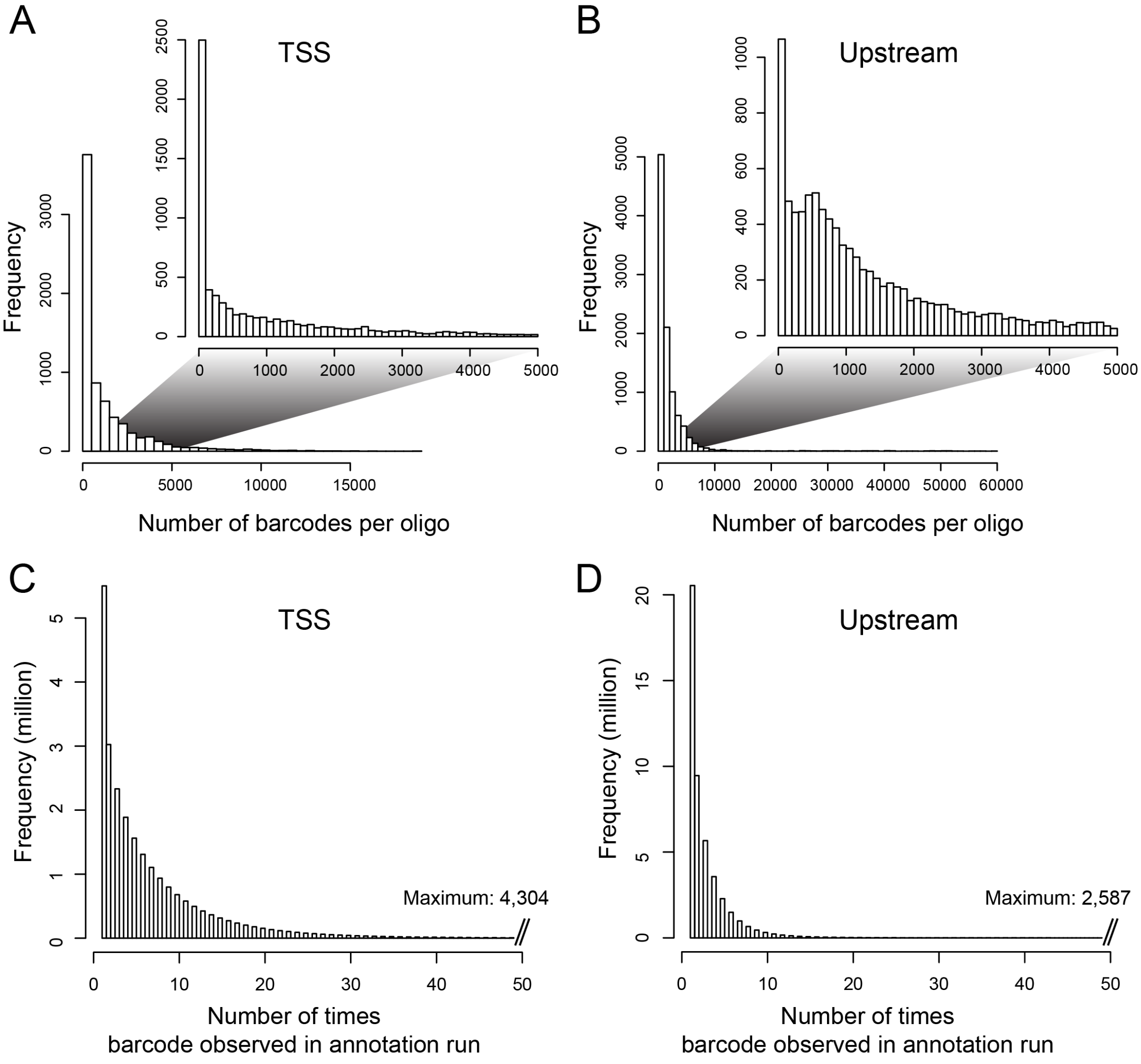


Figure S4: Distributions of barcodes. A) Number of barcodes tagging a given oligo in the TSS library. The inset shows the range from zero to 5,000 barcodes, which contains the majority of the distribution. B) as in A), but for the Upstream library. C) Distribution of the number of times a given barcode was observed in the TSS annotation sequencing run. D). As in C), but for the Upstream library.

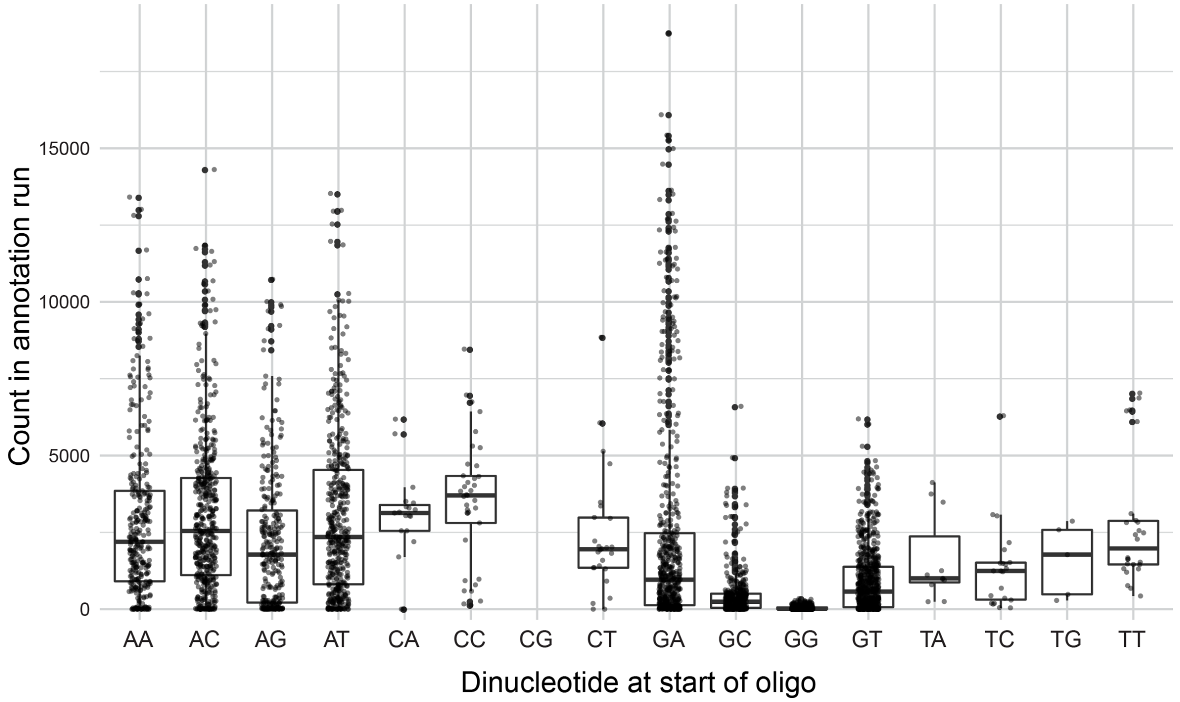


Figure S5: Number of times a given designed oligo was observed in the TSS annotation sequencing run as a function of the first two nucleotides of the oligo. Boxplots show the median as thick horizontal line, with the box showing the 25th and 75th percentiles. Whiskers show the largest value no further than 1.5 times the inter-quartile range; data points beyond this range are shown as individual dots. Note the reduced counts for oligos starting with a “G”, in particular those that started with a “GG”.

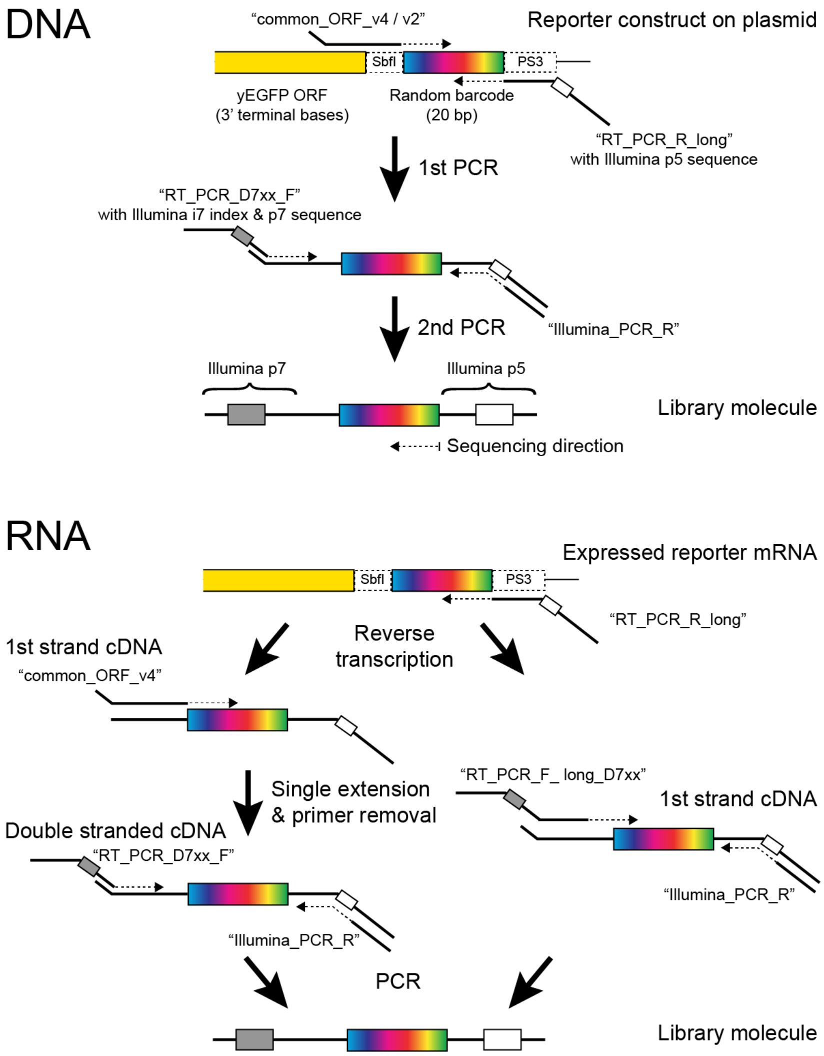


Figure S6: Barcode amplification. The figure shows details of the molecular reactions used to make Illumina sequencing libraries for barcode counting. Primer names are given in quotes. For RNA, Protocol 1 is on the left and Protocol 2 is on the right (see Methods for details). Note that in Protocol 1, the PCR step can exponentially amplify only cDNA but not plasmid molecules that may have escaped DNA degradation during RNA extraction because both PCR primers bind to overhangs added in the previous steps. If, during the single extension step, "common_ORF_v4" uses plasmid DNA as a template, the product lacks the p5 overhang, which contains the binding site for "Illumina_PCR_R". Conversely, if during single extension, "RT_PCR_R_long" (which is still present in the reaction) primes off a plasmid molecule, the product lacks the p7 overhang required by "RT_PCR_D7xx_F".

In protocol 2, the primers "common_ORF_v2" and "RT_PCRPD7xx_F" are replaced by primers "RT_PCR_F_long_D7xx". These primers permit direct amplification off plasmids and off first strand cDNA but require multiple long primers for multiplexing and provide less protection against inadvertent plasmid amplification.

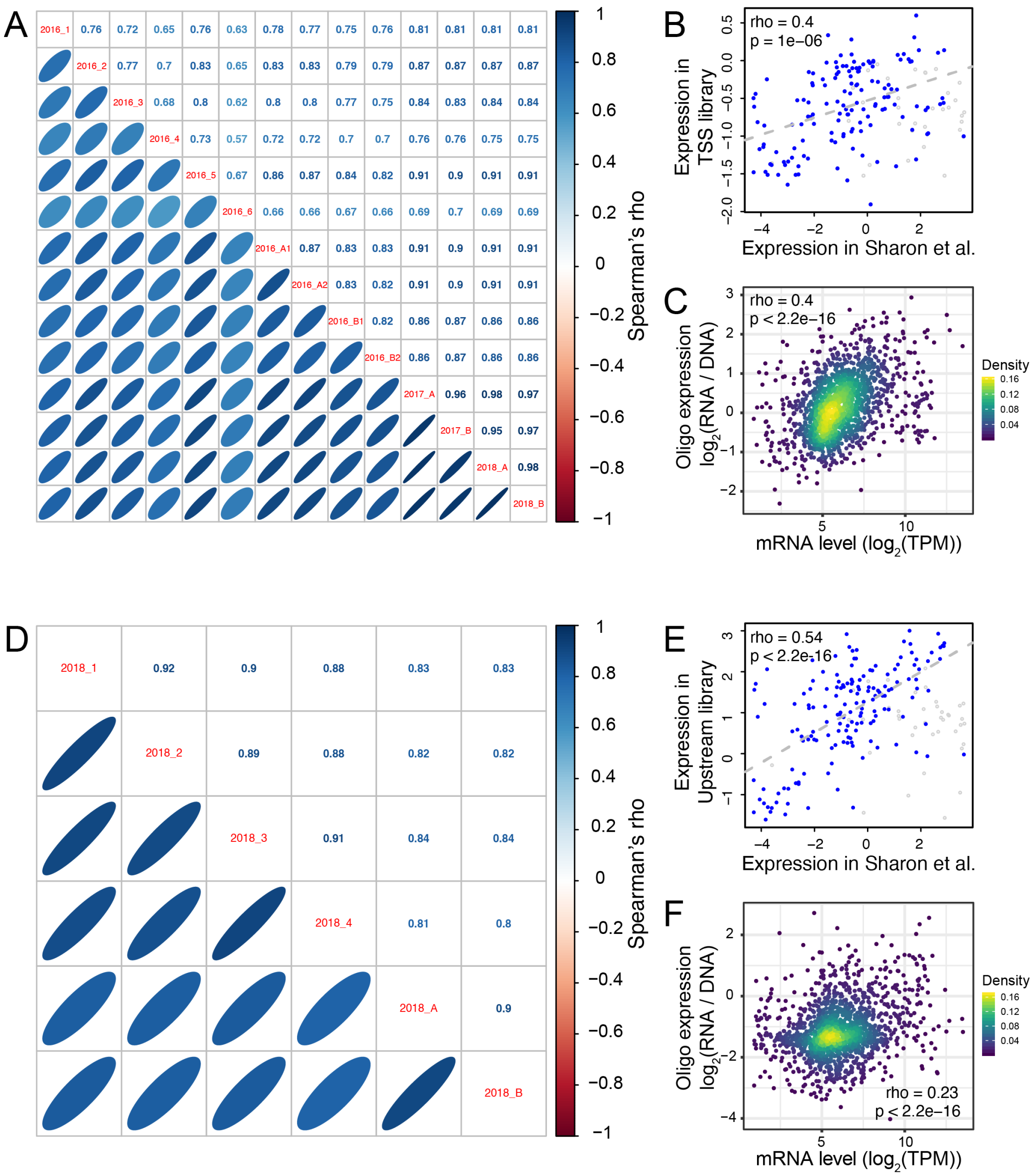


Figure S7: Reproducibility of oligo expression. A) Correlations of expression driven by oligos among replicates in the TSS library. B) Average expression across TSS replicates driven by the 200 oligos from Sharon *et al.*, (2012) compared to their published values. Blue points show oligos that did not include a Gal4 binding site; grey points show oligos that did include such a site. The indicated correlation was computed on oligos without Gal4 sites, because promoters with Gal4 sites are not expected to drive expression in the absence of galactose, as in the medium used here. Note that the correlation between observed and published data exists despite the different growth media between studies, and although Sharon *et al.* quantified expression by FACS-Seq instead of our RNA-Seq based measurements. C) Average expression driven by TSS oligos from a given gene promoter compared to that gene's mRNA level in the genome (Albert, Bloom, *et al.*, 2018). TPM: transcripts per million. D – F) as A – C, but for the Upstream library. The oligos in B & E showed higher correlations in the Upstream library than the TSS library, perhaps because the Upstream library but not the TSS library contained the same minimal *HIS3* promoter fragment used in Sharon *et al*. The correlation with native genes (C & F) was stronger in the TSS than the Upstream library, perhaps because TSS oligos more closely resembled a native promoter due to their closer proximity to the transcription start sites and the TSS library's lack of the minimal *HIS3* promoter.

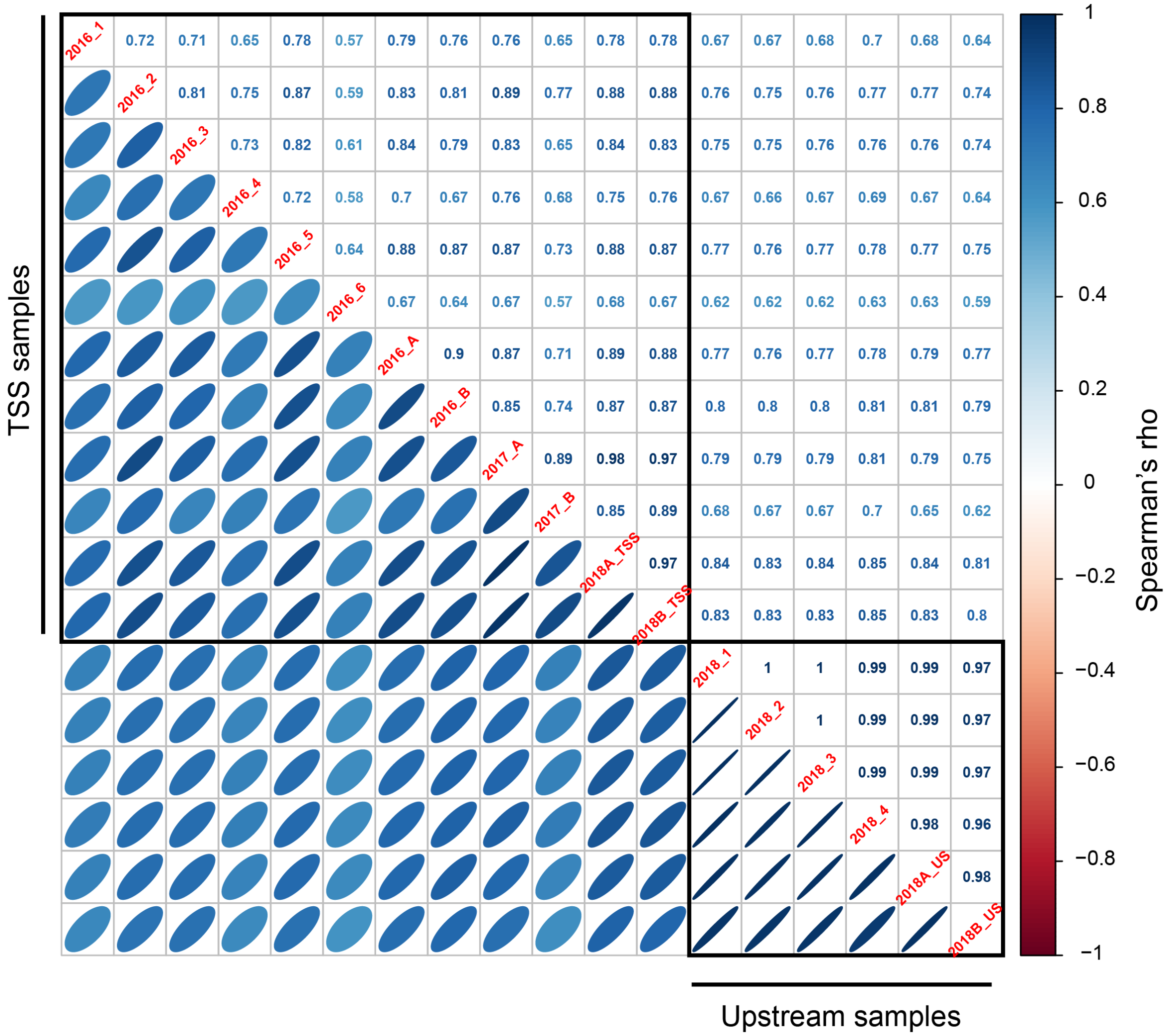


Figure S8: Correlations of expression driven by the 200 oligos common to the TSS and Upstream library.

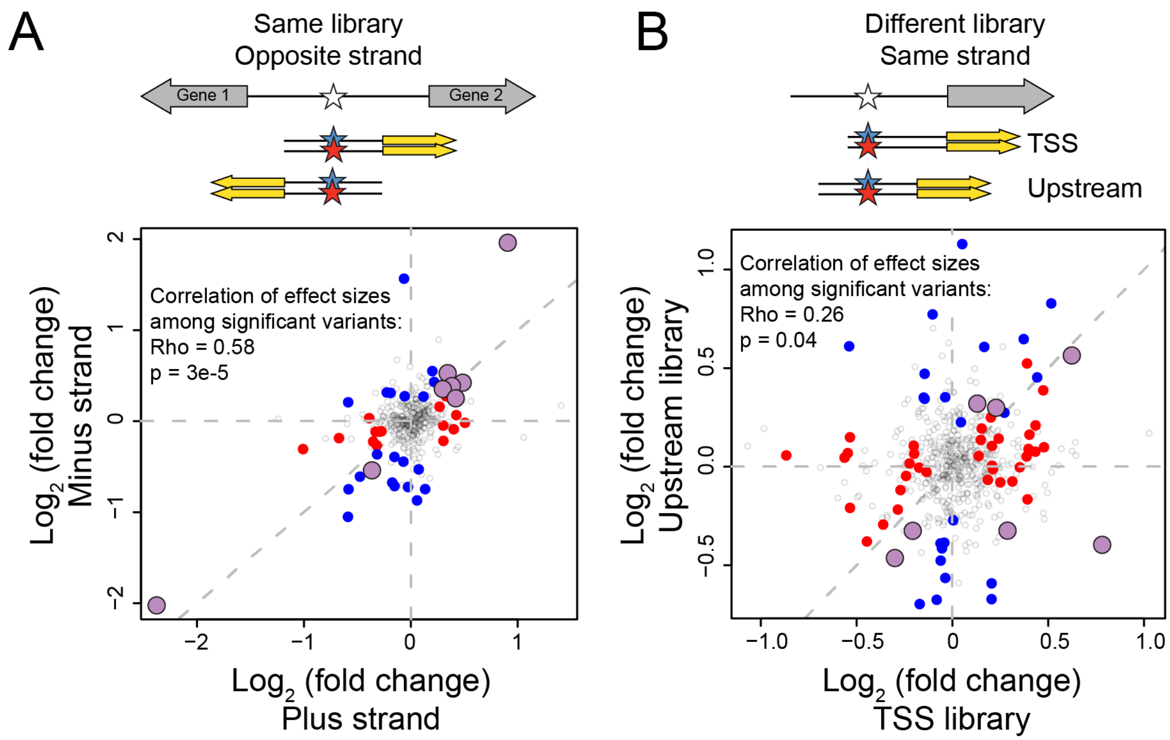


Figure S9: Reproducibility of variant effects. A) Variants measured in the same library (TSS or Upstream) but in opposite orientation ("strand") with respect to the reporter gene. Blue points: variants with significant (5% FDR) effects on the plus strand. Red points: variants with significant effects on the minus strand. Purple, larger points: variants significant in both orientations. The indicated correlations were computed for variants that were significant in at least one of the two orientations. B) As in A) but comparing effects of variants measured on the same strand but in the two different libraries.

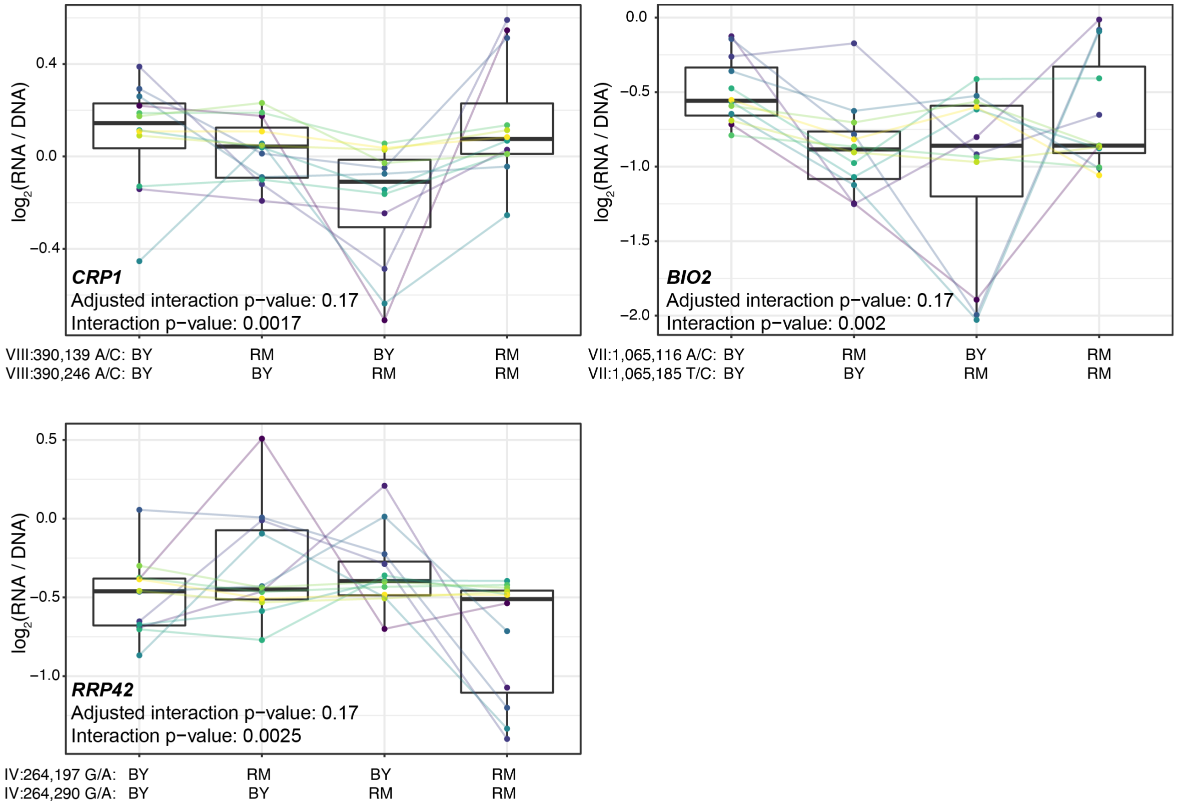


Figure S10: Additional cases of significant epistasis between promoter variants. Each panel shows, for one gene, MPRA expression driven by four oligos with the indicated combination of BY and RM alleles at the two variants. Each panel states the gene name in bold along with multiple-testing adjusted as well as raw interaction p-values. Variants are given as “chromosome:position reference/alternative allele”. Colored lines between boxplots connect the data for a given oligo in the different biological replicates. Boxplots show the median as thick line, with the box showing the 25th and 75th percentiles. Whiskers show the largest value no further than 1.5 times the inter-quartile range; any observations beyond this range are shown as individual points.
